# Evolution in Eggs and Phases: experimental evolution of fecundity and reproductive timing in *Caenorhabditis elegans*

**DOI:** 10.1101/042143

**Authors:** Bradly Alicea

## Abstract

To examine the role of natural selection on fecundity in a variety of *Caenorhabditis elegans* genetic backgrounds, we used an experimental evolution protocol to evolve 14 distinct genetic strains over 15-20 generations. Beginning with three founder worms for each strain, we were able to generate 790 distinct genealogies, which provided information on both the effects of natural selection and the evolvability of each strain. Among these genotypes are a wildtype (N2) and a collection of mutants with targeted mutations in the *daf-c*, *daf-d*, and AMPK pathways. The overarching goal of our analysis is two-fold: to observe differences in reproductive fitness and observe related changes in reproductive timing. This yields two outcomes. The first is that the majority of selective effects on fecundity occur during the first few generations of evolution, while the negative selection for reproductive timing occurs on longer timescales. The second finding reveals that positive selection on fecundity results in positive and negative selection on reproductive timing, both of which are strain-dependent. Using a derivative of population size per generation called the reproductive carry-over (RCO) measure, it is found that the fluctuation and shape of the probability distribution may be informative in terms of developmental selection. While these consist of general patterns that transcend mutations in a specific gene, changes in the RCO measure may nevertheless be products of selection. In conclusion, we discuss the broader implications of these findings, particularly in the context of genotype-fitness maps and the role of uncharacterized mutations in individual variation and evolvability.

## INTRODUCTION

There is a rich tradition of using the Nematode *Caenorhabditis elegans* to make controlled genetic manipulations that provide significant insights regarding genetic effects on development and physiology. Experimental Evolution methods provide us with the opportunity to look beyond the classical genetic methods [1], and provides us with a means to study the effects of natural selection in a way not possible with comparative or inferential techniques [2]. In particular, experimental evolution allows control and temporal resolution over the evolutionary process, which enables novel functional assessments of existing mutant genotypes [3]. Using a dataset of a single wildtype genotype and 13 mutant genotypes, we will address three interrelated questions:

A) What effect does natural selection and mutational diversity have on fecundity (as measured by population size over a finite interval)?
B) Are the observed differences in population size informative with respect to genotypic identity?
C) Do we observe changes in reproductive timing (as measured by Reproductive Carry-Over) that result from positive selection for fecundity?

We have generated between 47 and 64 genealogies per strain for 14 genetic strains of *C. elegans* over 15-20 generations (see Table 1, full information in Supplemental Data). These genealogies result from repeated selection during each generational interval (3d). We terminate all non-selected replicates in every generation. Founder worms at the L4 stage of development selected in terms of largest population sizes generated after 3d. Founder worms for the next generation are drawn randomly from the selected populations at proportions consistent with the relative size of the selected populations. This process produces a series of new replicates representing the subsequent generation. In this way, we maintain the same number of replicates during every generation in the experiment. As we repeatedly select some populations over others, we generate a large number of genealogies of varying length.

**Table 1.**
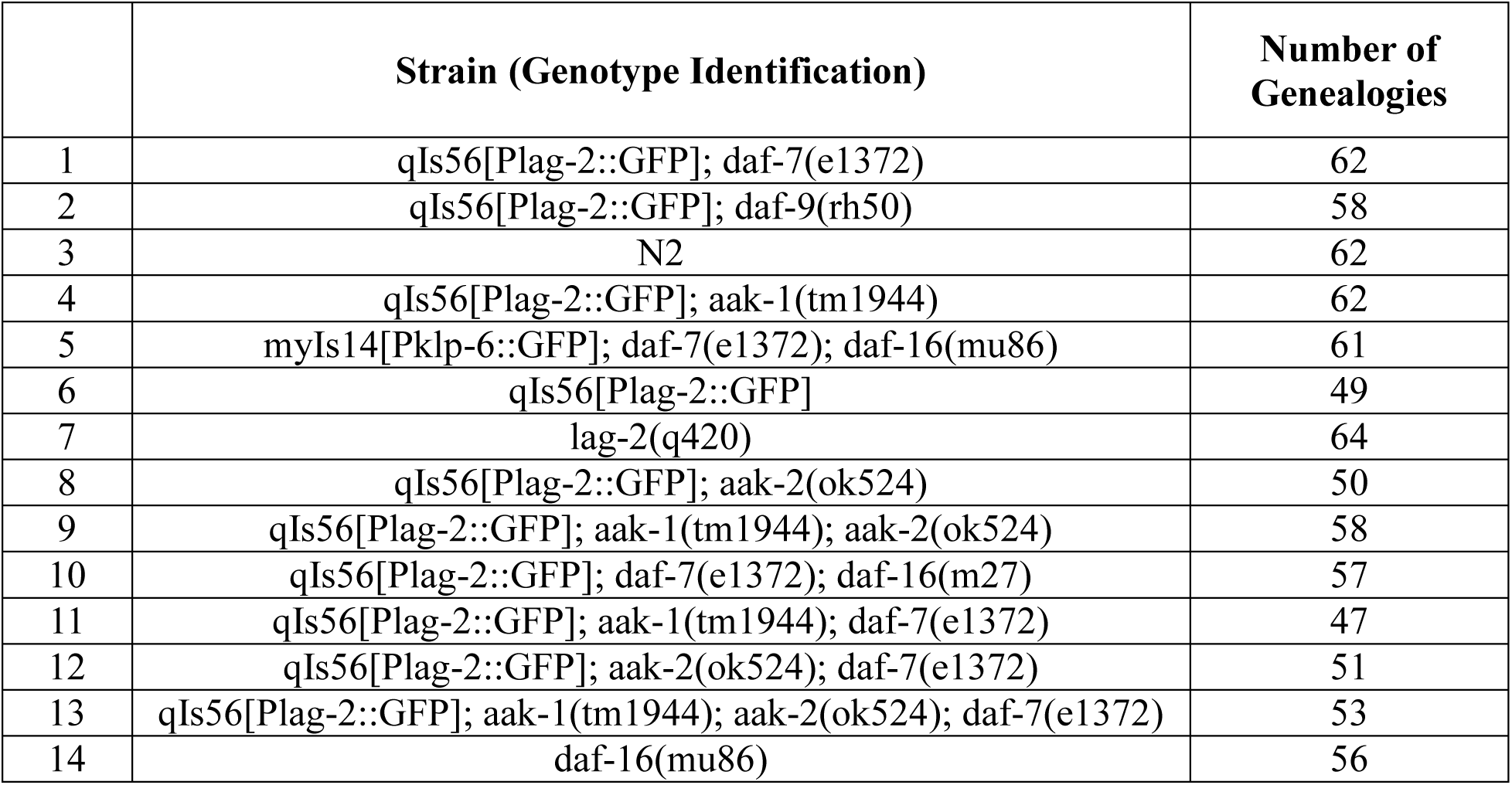
An inventory of strains and the number of genealogies generated via natural selection.

In this study, we are not only selecting for fecundity, but in some cases also indirectly selecting for reproductive timing. While there are a finite number of eggs and sperm in each individual worm, their reproductive capacity still exhibits some variability. Some of this variability is stochastic in nature, but a major component is strongly influenced by either environment or genetic background. All things being equal in terms of environment, selecting from populations with large numbers of offspring 3d after the founder worm's L4 stage will favor early reproducers in subsequent populations. By keeping the environment fixed across the course of our evolutionary trajectories, it is our goal to demonstrate the selective effects across a diverse range of mutant genotypes.

According to Gray and Cutter [4], life-history and the effects of mutation are a fertile area for *C. elegans* experimental evolution research. In this study, we intend to take advantage of this by examining the evolvability of a variety of genetic mutant strains when positively selected for fecundity in earlier portions of the reproductive cycle. Selecting for fecundity often results in evolutionary changes over a relatively low number of generations because reproduction is sperm-limited [5]. In wildtype genetic backgrounds, life-history related shifts in sperm production [6] can lead positive selection for fecundity. In a previous study [7], it was shown that wildtype isolates can be selected for earlier reproduction after 47 generations of evolution. In addition, this positive selection is also be decoupled from decreases in lifespan and later lifespan fecundity.

The ability to impose positive selection for fecundity on populations over relatively short timespans also suggests that experimental evolution can uncover new pathways to the evolution of complex traits. This includes influencing the evolvability and robustness of the genealogies that result from sustained natural selection. In [8], the Nematode species *Caenorhabditis remanei* underwent natural selection for heat shock resistance over 10 generations. This form of environmental selection results in negative selection for robustness to heat shock. A less robust response meant that descendent generations were much less resistant to heat shock than non-selected organisms. Yet selection may not be the key driver in such interactions. According to constructive neutral theory [9], mutation is the primary driver of constraints, compensatory functions, and novelty, while selection acts merely to filter this variation in various ways. We can observed this to some extent in the form of restoring fitness advantages lost to mutational drift. For example, compensatory mutations play a role in restoring fitness amongst individual worms in mutation-accumulation lines of *C. elegans* [10] when being derived from large populations and undergoing natural selection [11].

## RESULTS

The first question involves examining the effects of selection and mutational diversity on fecundity. To do this, we will first establish the mean level of fecundity within specific genetic strains. Then, we will demonstrate the effects of selection for fecundity by comparing mean population sizes across strains. Using a statistical analysis called exponential smoothing (Figure 1), we can derive a population mean per generation for 14 strains described in Table 1 that removes many of the demographic fluctuations observed in the raw data (Figure 1).

**Figure 1.**
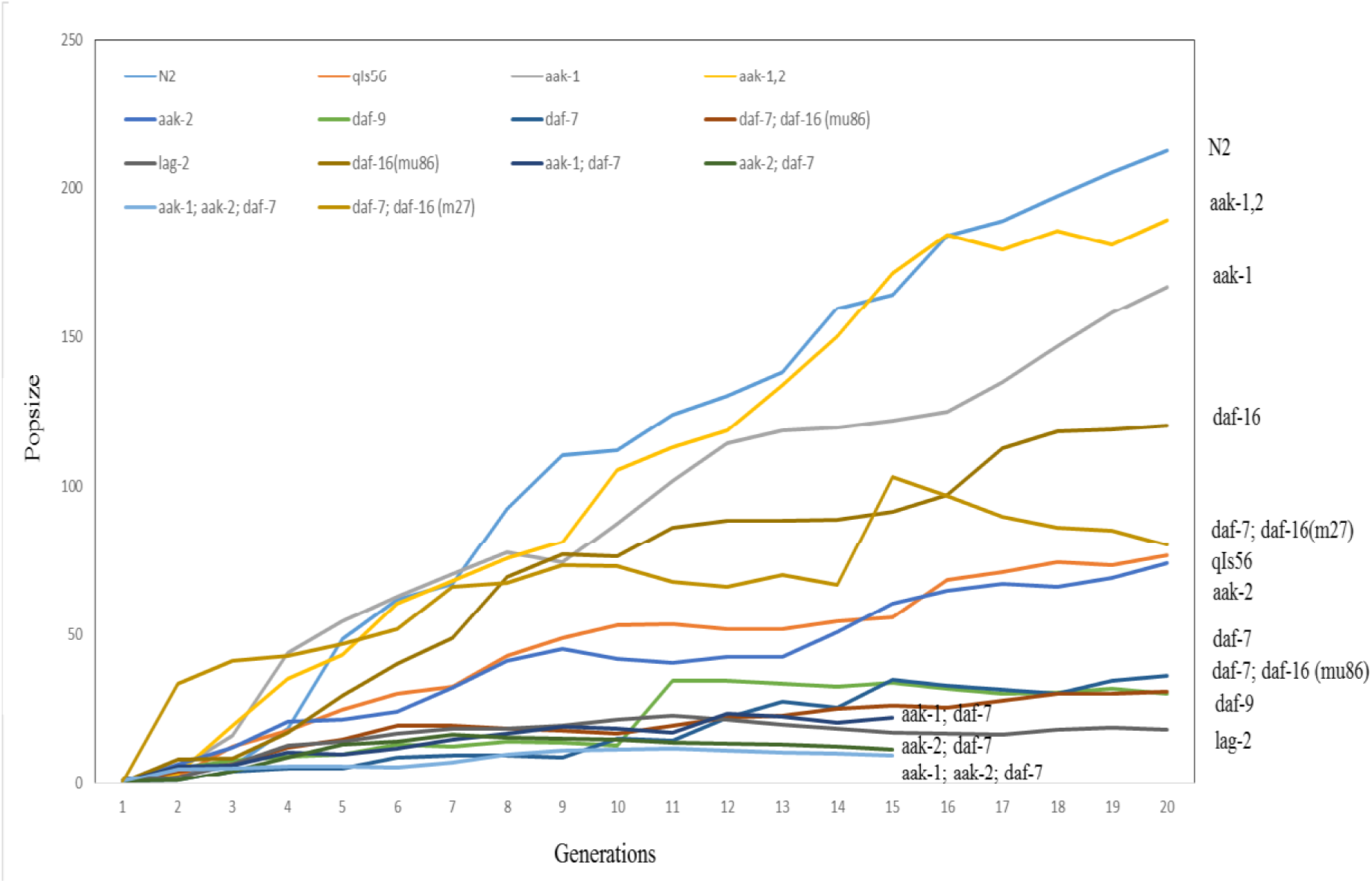
Evolutionary trajectory (measured using mean population size per generation) for 11 strains over 20 generations and 3 strains over 15 generations, normalized using exponential smoothing (α = 0.1).

According to the analysis in Figure 1, we should expect the wildtype (N2) to exhibit the greatest increase in fecundity, followed by the two AMPK mutants aak-1(tm1944) and aak-1(tm1944); aak-2(ok524), and then the daf-16(mu86) mutant. Notice that the *daf-c* and *lag* mutants exhibit the least pronounced gains in fecundity. In Figures 2 through 5, we will look at a slightly different characterization of mean population size (mean population size for every genealogy) to examine fitness gains relative to fecundity.

**Figure 2.**
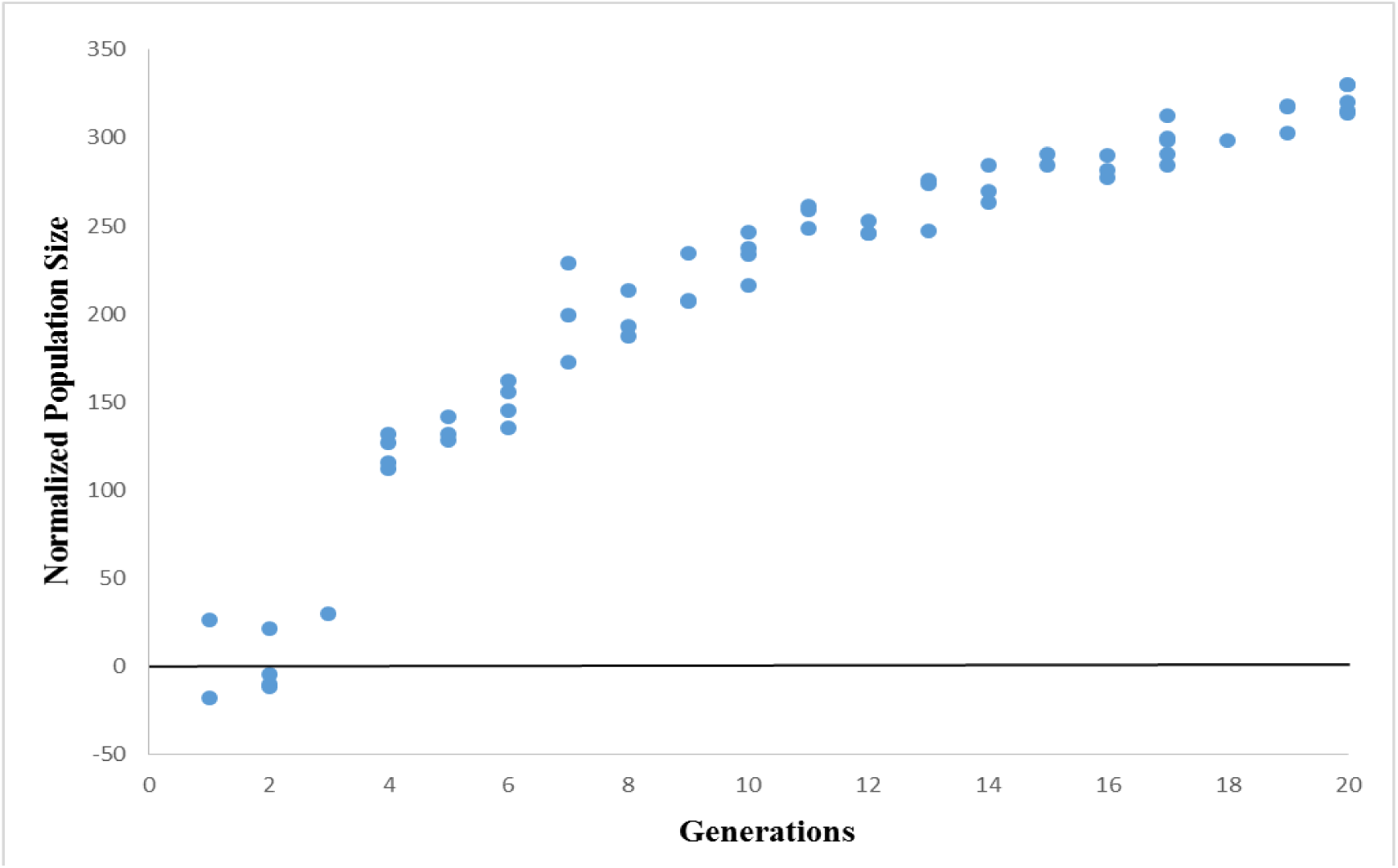
20 generations of experimental evolution on the wildtype (N2) strain (62 geneaologies). Normalized population size is a non-evolved control subtracted from the measured population size.

We can also examine fecundity gains by comparing the evolved population size with an unevolved population size measurement (see Methods, Normalized Population Size). Figure 2 shows that for the wildtype (N2), there is an initial period (three generations) where the evolved population size is roughly comparable to that of the unevolved population size. However, beginning at generation 4, there is a jump in normalized population size that progresses logarithmically until generation 20. While the logarithmic signature is expected due to sampling bias across the length of a genealogy, this demonstrates a signature of positive selection for sustained selective pressure on early fecundity.

Figure 3 shows two sets of outcomes for the AMPK mutants (Figure 3). For the aak-2(ok524) mutant, there is a jump in normalized population size at Generation 2 which plateaus at a value of 100 after Generation 10. In the case of the aak-1(tm1944) and aak-1(tm1944); aak-2(ok524) mutants, there are two jumps in normalized population size before Generation 5. This leads to a logarithmic increase as they approach Generation 20. At Generation 20, both the single aak-1 and double mutant exhibit normalized population size values comparable to that of N2.

**Figure 3.**
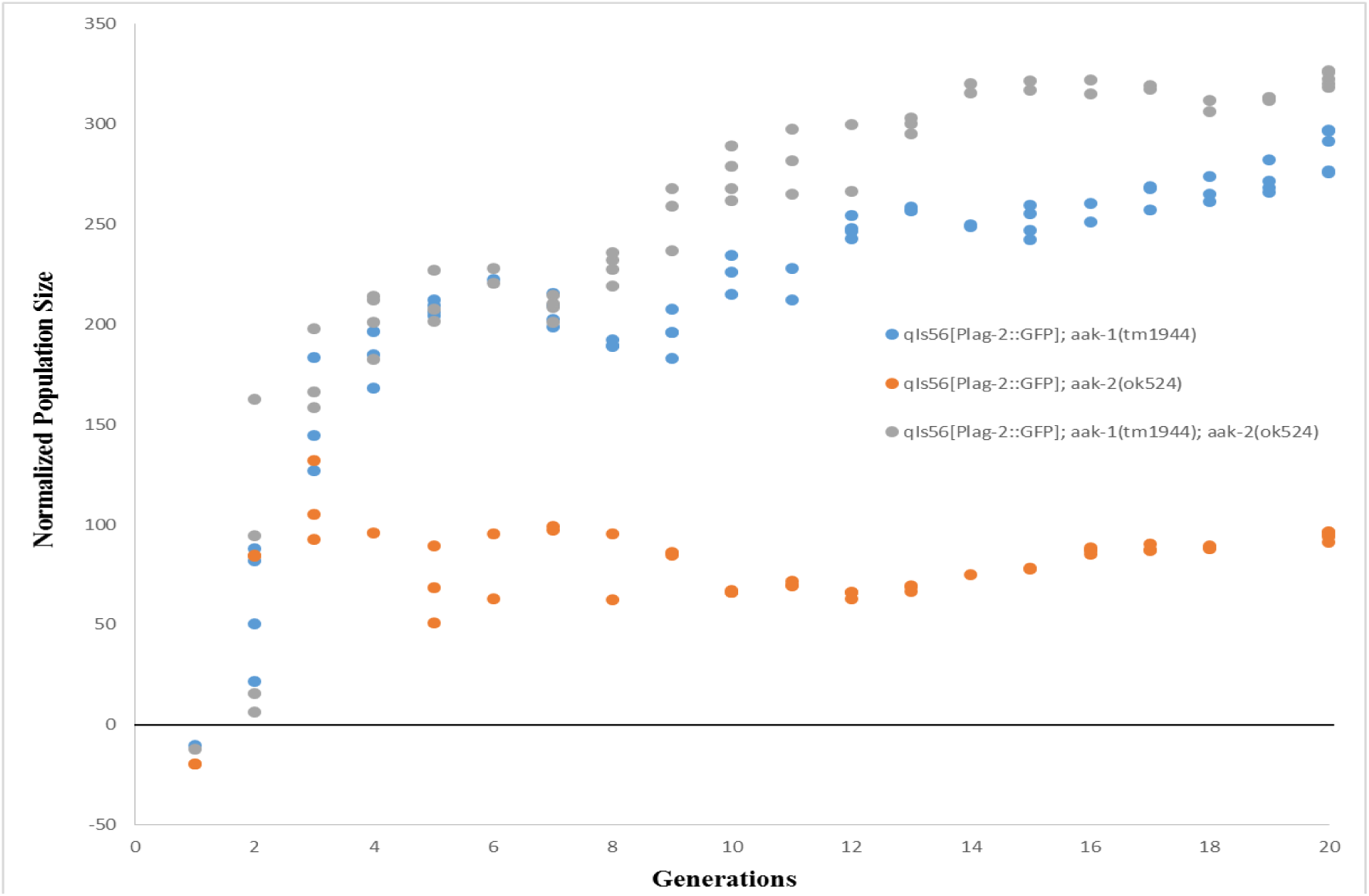
20 generations of experimental evolution on the AMPK mutant strains (62 genealogies for *aak-1*, 50 genealogies for *aak-2*, and 58 genealogies for *aak-1; aak-2*). Normalized population size is a non-evolved control subtracted from the measured population size.

In the case of the daf-7(e1372) mutants (Figure 4), there is the same logarithmic progression of fecundity observed in Figures 2 and 3, but originating at a much lower starting point. As is also the case with Figures 2 and 3, the daf-7(e1372) mutants exhibit an early spike (at Generation 2) in fecundity. Figure 5 shows the normalized population size over 15 generations for the three AMPK/daf-7(e1372) mutant genotypes. As with the AMPK and daf-7 mutant genotypes described in Figures 3 and 4, we observe an early spike in fecundity culminating in peak fecundity during Generation 6 for the single mutants aak-1(tm1944); daf-7(e1372) and aak-2(ok524); daf-7(e1372). For longer genealogies, however, we do not see the same logarithmic increase. Instead, we observe a decline in fecundity and intra-strain variation after Generation 10.

**Figure 4.**
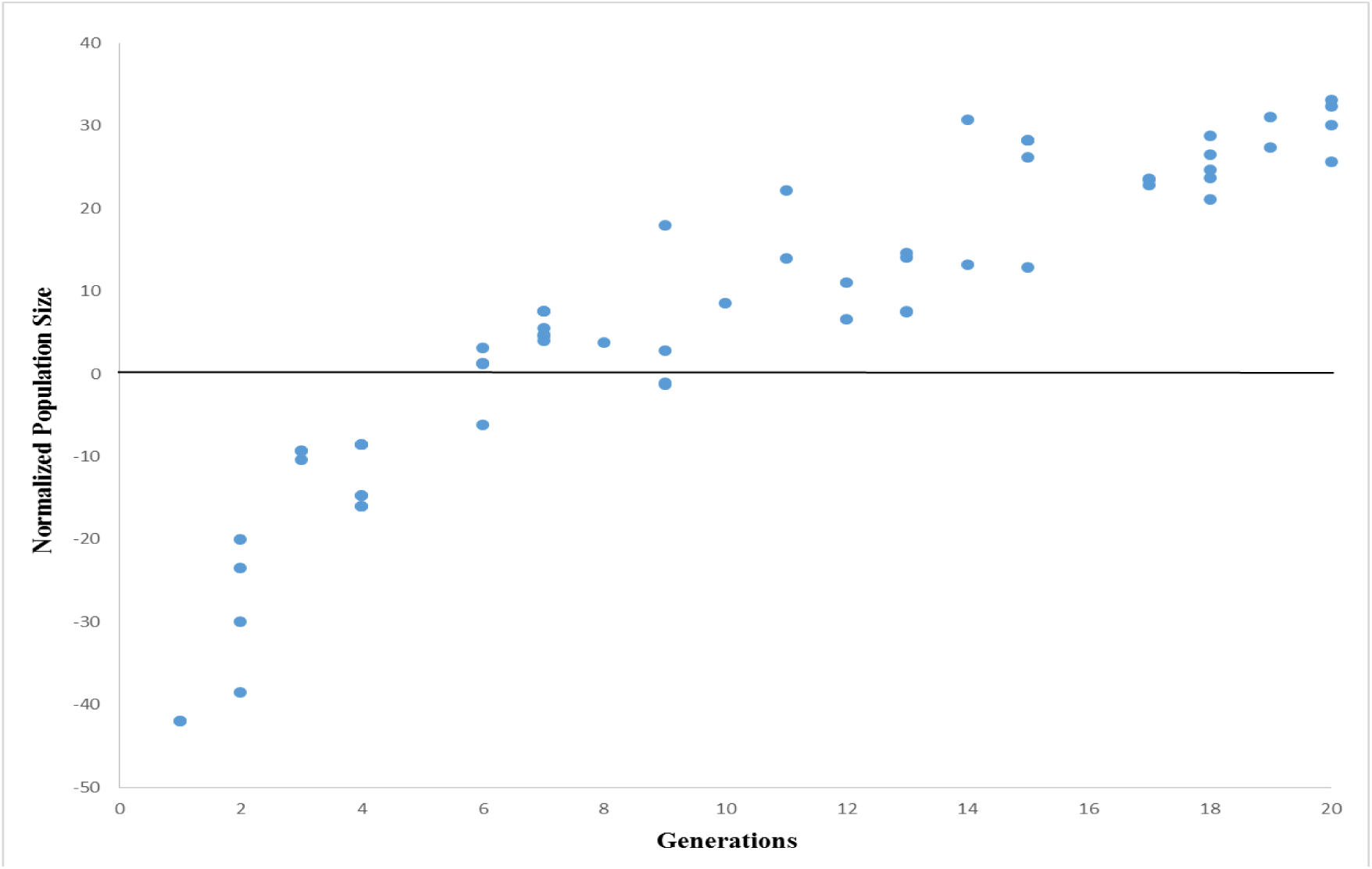
20 generations of experimental evolution on the daf-7(el372) strain (62 genealogies). Normalized population size is a non-evolved control subtracted from the measured population size.

**Figure 5.**
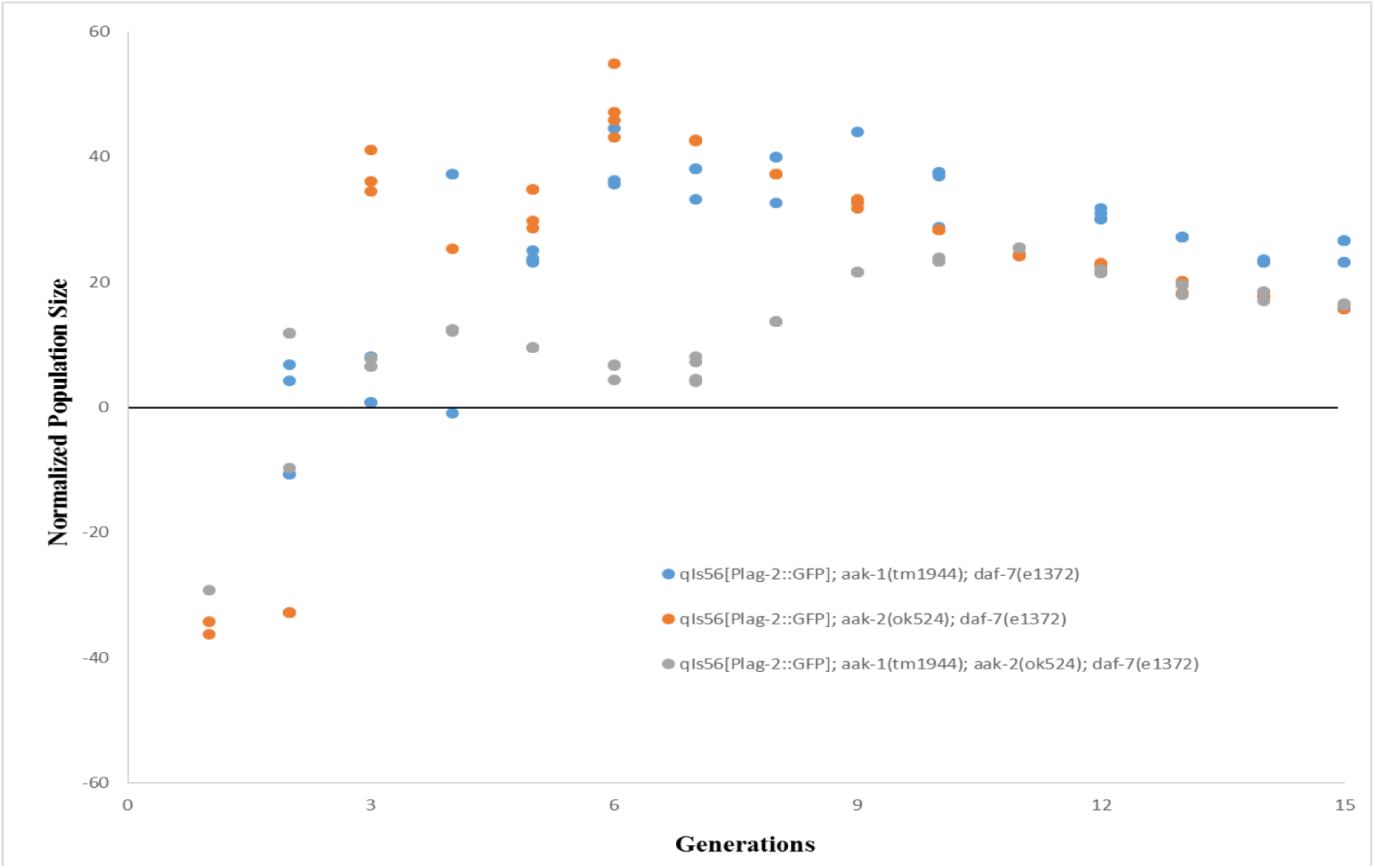
15 generations of experimental evolution on the AMPK/*daf-c* mutant strains (47 genealogies for aak-1; daf-7, 51 genealogies for aak-2; daf-7, and 53 genealogies for aak-1; aak-2; daf-7). Normalized population size is a non-evolved control subtracted from the measured population size.

The second question can be answered with a more complex analysis of the population size. In wild isogenic lines, Diaz and Viney [12] have observed inter-line differences in mean reproductive variance that are negatively related to mean lifetime fecundity. In this context, we can look at the variance per generation across genealogies by calculating the Reproductive Carry-over (RCO). RCO is a first-order derivative of the population size genealogies. Figure 6 shows a heat map for the distribution of RCO values across the evolutionary trajectory for 14 strains.

**Figure 6.**
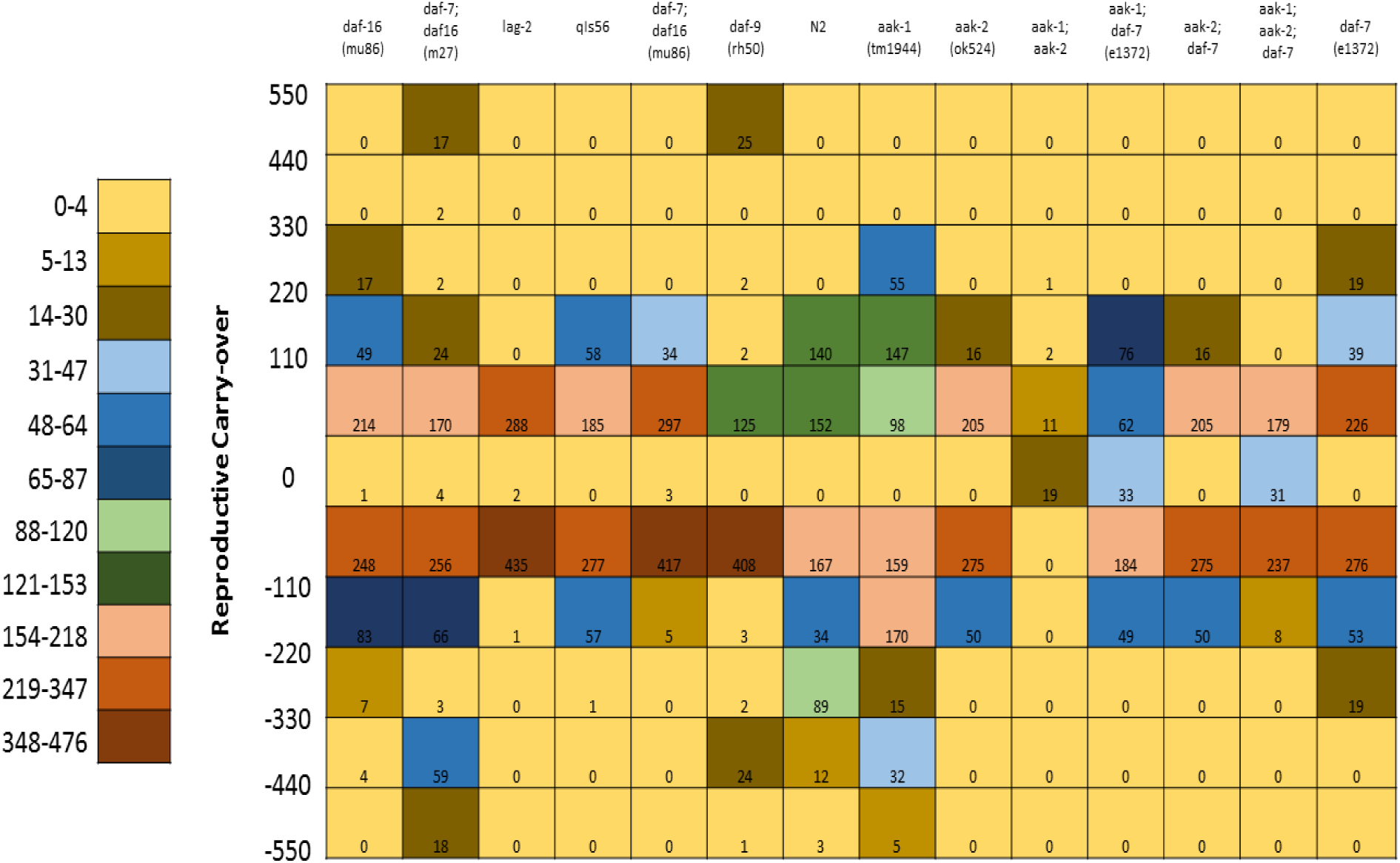
A heat map of Reproductive Carry-Over (RCO) measurement values for 14 strains (13 mutant strains and 1 wildtype) over 15-20 generations. All bins (vertical axis) are of size 110 except for the zero bin.

These data reveal a number of inter-strain differences not directly linked to mutations in specific genes. To further appreciate this, we can compare Figures 1 and 6, which demonstrate the smoothed population size over evolutionary time and the distribution of RCO values, respectively. Strains such as *lag-2*, *aak-2*, and the AMPK/*daf-7* mutants do not attain large population sizes over the course of their evolutionary trajectory. As a result, they also do not exhibit many negative RCO values. In other strains that never attain large population sizes (*daf-7* and *daf-9*), negative RCO values can be tied to demographic fluctuations (or, more specifically, the downward portion of that fluctuation).

A more subtle effect involves evidence for positive selection on developmental delay. In some strains, as individual worms were positively selected for fecundity, there was corresponding negative selection on developmental timing. For example, in the AMPK/daf-7 mutants, the 3d reproductive period that defined a generation was often only enough time for the adult worm to lay eggs. By contrast, in strains such as N2 the 3d reproductive period that defined a generation produced many L4 stage offspring to select from. This difference can be seen quantitatively in Figure 6 in that strains that exhibited this phenomenon also tended to have RCO values that were strongly negative or exhibit RCO values that range from 150 to ‐150.

To compare the distribution of RCO values between selected strains in terms of their probability distribution, we plot the Cumulative Distribution Function (CDF) for each strain and compare their shape. This can give us more information about the statistical context of RCO measurements for each strain. Figure 7 shows a comparison between the wildtype (N2), a *daf-c* control (*daf-7*), and three AMPK mutants (*aak-1*, *aak-2*, and *aak-1*; *aak-2*). The wildtype (N2) and *aak-1* strains have similar distributions, as do the *daf-c* control (*daf-7*) and *aak-2*. The double mutant (*aak-1*; *aak-2*) shares features of both pairs. All strains differ very little around the mean, which is observed in the heat map as well.

**Figure 7.**
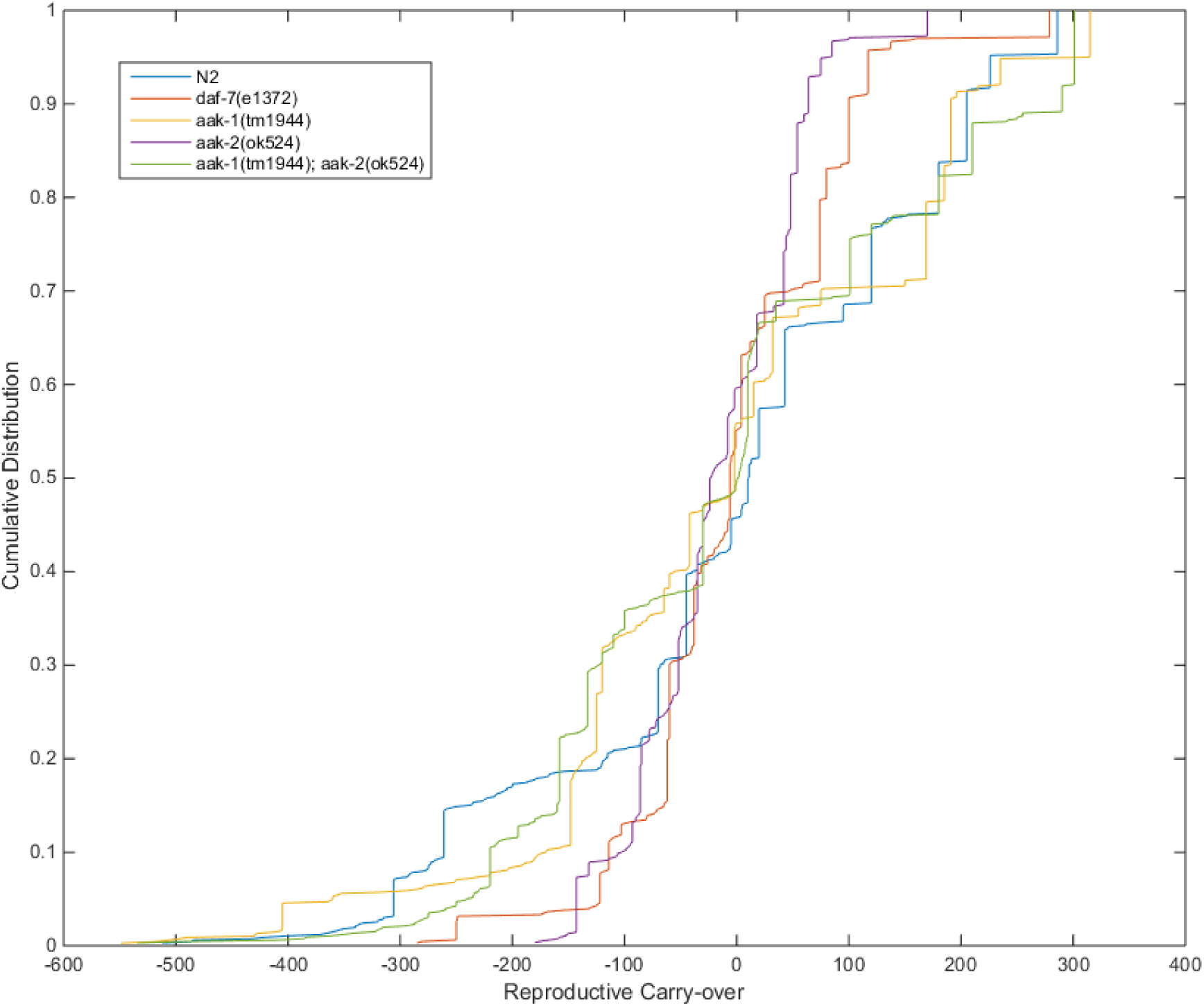
Cumulative distribution functions (CDFs) of the RCO measure for three AMPK mutant genotypes, a *daf-c* mutant (*daf-7*), and the wildtype (N2).

Figure 8 shows data for the wildtype (N2) and daf-c control (daf-7) shown in Figure 7, in addition to the three AMPK/daf-7 strains (*aak-1;* daf-7, *aak-2; daf-7*, and *aak-1; aak-2; daf-7*). In this case, the CDF for N2 is quite distinct from the CDFs for the *daf-7* and AMPK/daf-7 genetic backgrounds. In particular, N2 differs from the other strains in terms of the tails of its distribution. By contrast, the *aak-2; daf-7* and *aak-1; aak-2; daf-7* strains exhibit extremely short tails, which underscores the lack of variation and overall lack of demographic fluctuation in these genotypes.

**Figure 8.**
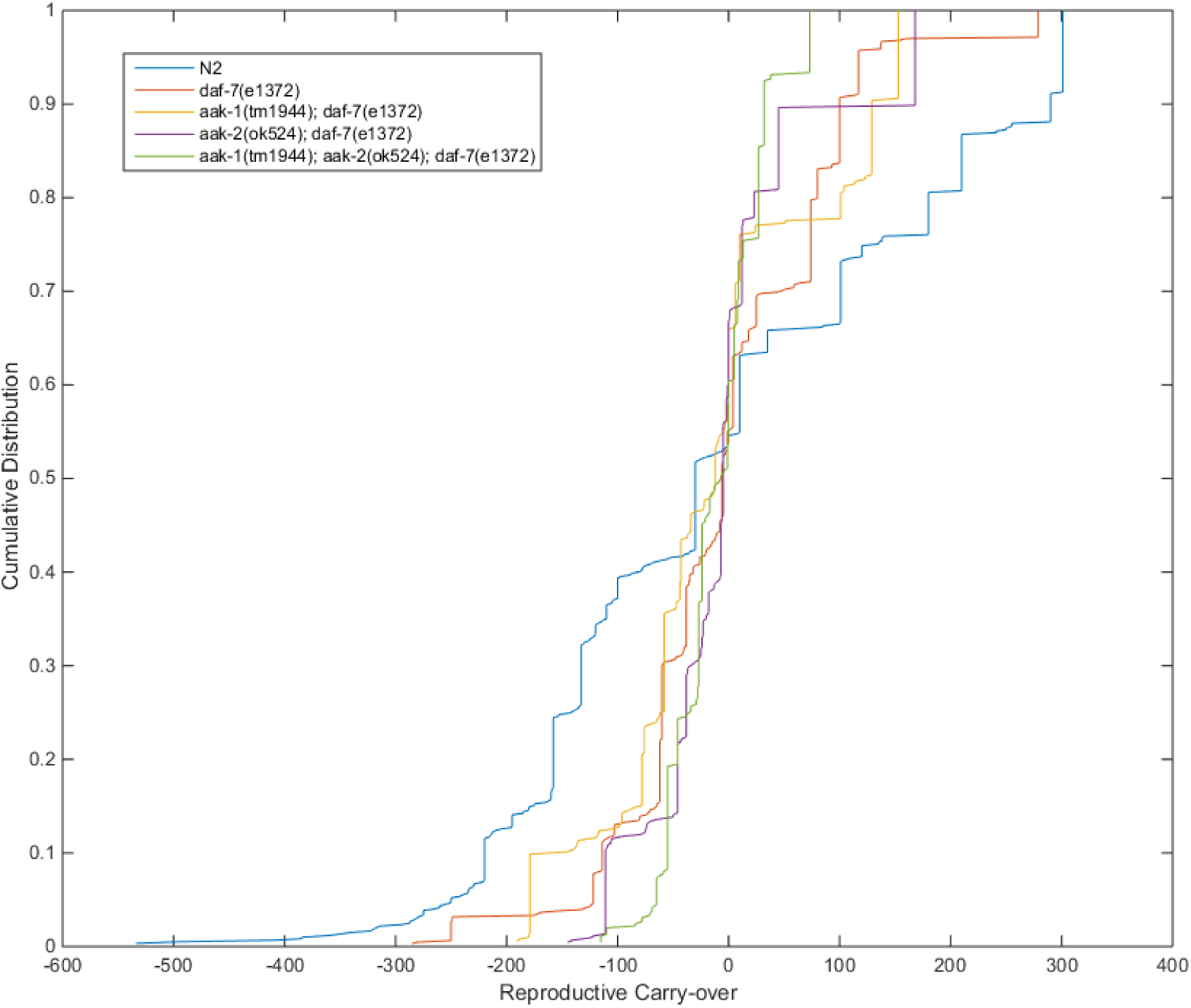
Cumulative distribution functions (CDFs) of the RCO measure for three AMPK/daf-c mutants (aak-1; daf-7, aak-2; daf-7, and aak-1; aak-2; daf-7) mutant genotypes, a daf-c mutant (daf-7), and the wildtype (N2).

To demonstrate that there is in fact selection for reproductive timing, we must make a more implicit connection between reproductive potential and the presence of offspring themselves. Our third question is answerable by identifying potential signatures of selection in the evolutionary data. We accomplish this by sampling an evolutionary time series every 4-5 points and comparing population size and its standard error. In Figure 9, we can see the normalized population size at five different points within a 20 generation series for the wildtype (N2) and three different AMPK mutant strains (*aak-1(tm1944)*, *aak-2(ok524)*, *aak-1(tm1944)*; *aak-2(ok524))*. These results confirm some of the outcomes observes in Figures 1 and 2. For example, in the wildtype (N2) and two mutant genotypes (*aak-1*, *aak-1; aak-2*), there is an early shift towards larger mean population sizes before Generation 5 (Figure 8, A, B, and D).

**Figure 9.**
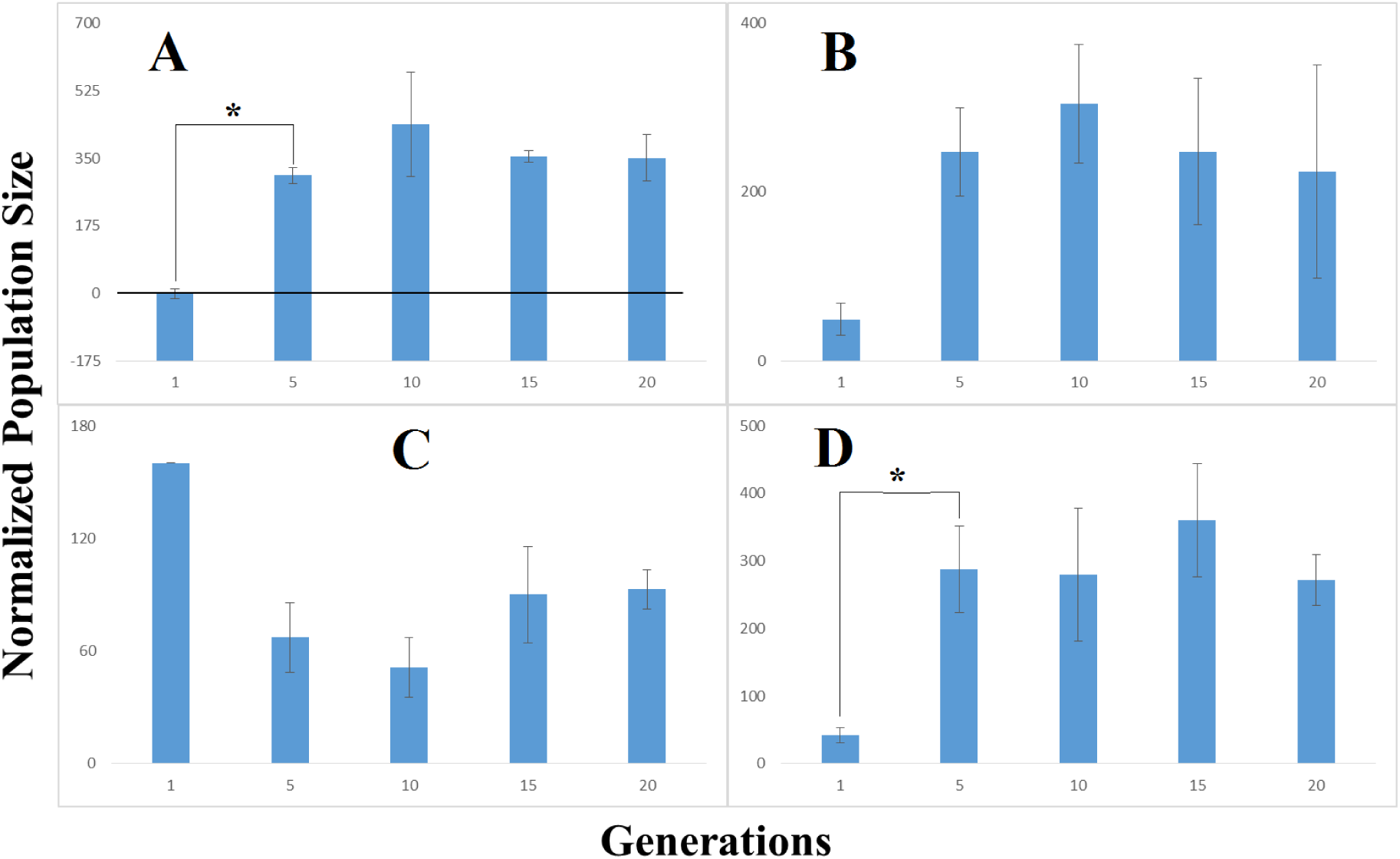
Normalized population size versus five discrete generations of the wildtype (N2) genotype and three AMPK mutant genotypes. Clockwise from upper left: N2 (A), aak-1 (B), aak-2 (C), aak-1; aak-2 (D). Starred pairs are statistically significant (two-tailed t-test).

Based on the results of a Bonferroni-corrected t-test, the shifts for N2 and *aak-1; aak-2* between Generations 1 and 5 are statistically significant, p<0.03 and p<0.02, respectively. There seems to be a lack of selection afterward, although the standard errors for the *aak-1* mutant genotype (Figure 9, B) become more pronounced in later generations. In the case of the *aak-2* mutant genotype (Figure 9, C), we see an opposite pattern from the other AMPK mutants. After an initial dropoff in fecundity that is not statistically significant, there appears to be a continued lack of selection. This is distinct from negative selection for fecundity, as the population size measure does not produce any statistically significant decreases.

Figure 10 shows a more mixed result. In this comparison, the daf-7(e1372) strain is used as a control (Figure 10, A). This shows a large increase normalized population size at Generation 5, accompanied by a dropoff in normalized population size for Generations 10 and 15. Normalized population size increases once again for Generation 20. Conservatively, we can say that this increase is due to demographic fluctuation, and is not due to quick evolutionary changes. This is confirmed by the reproductive behavior of the AMPK/daf-7 mutants (Figure 10, B-D), all of which exhibit transient fluctuations. The only strain to show some evidence of negative selection of normalized population size over time is aak-1(tm1944); daf-7(e1372). In this strain (Figure 10, B), normalized population size is above the control in Generation 1 and 5. Somewhere between Generation 5 and Generation 10, normalized population size drops below the control and remains there for the rest of the experiment.

**Figure 10.**
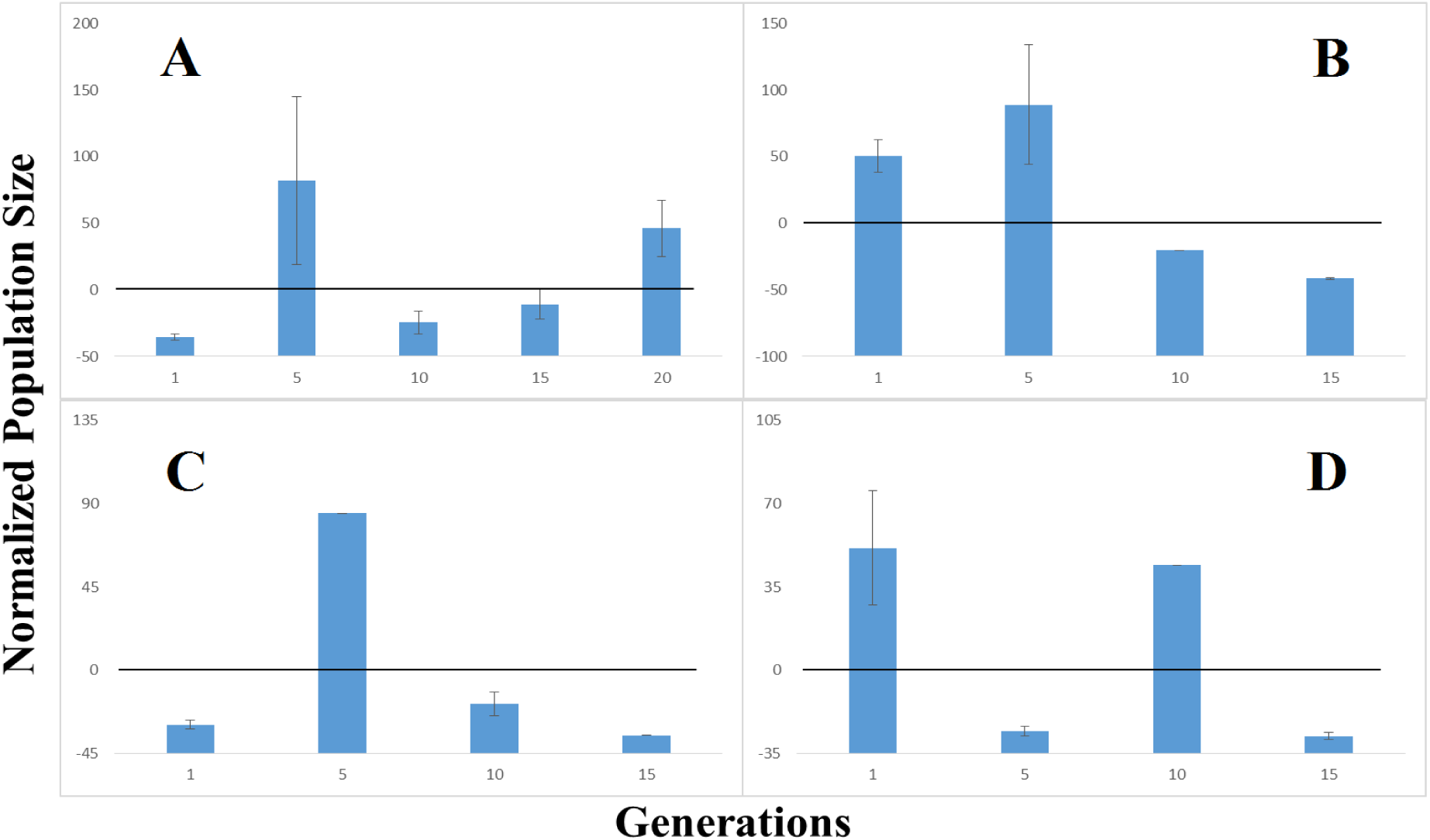
Normalized population size versus five discrete generations of the daf-7 mutant genotype and three AMPK/daf-c mutant genotypes. Clockwise from upper left: daf-7 (A), aak-1; daf-7 (B), aak-2; daf-7 (C), aak-1; aak-2; daf-7 (D).

The effects of positive selection on fecundity appear to be strain-dependent. For our wildtype strain, positive selection did indeed lead to greater fecundity. In this case, selecting for larger brood sizes did lead to a speed-up in reproductive timing whereby larger brood sizes were maintained. However, in the case of our daf-c and AMPK/daf-c mutant strains, positive selection for greater fecundity actually lead to smaller brood sizes over our generational interval. This may be due to some form of reproductive delay, which in the case of our AMPK/daf-c mutant strains drives our genealogies to premature extinction.

## DISCUSSION

Taking these data collectively, we can see that there are two effects of positive selection on fecundity. These differing effects are contingent upon the genetic background of the mutant strain in question. One effect is positive selection on reproductive timing, which results in nonlinear population size increases. This is observed in the wildtype, in addition to the AMPK mutants. The alternative effect is negative selection on reproductive timing, which results in no clear signature of selection on population size and minimization of both the Smoothed Population Size and the RCO measure.

We can also see a clear nonlinearity in terms of selective advantage. In a number of strains, there appears to be an early phase and a later phase of evolution. In the early phase, we see either a large and unambiguous jump in fecundity along with large variability in the RCO. Characteristics of later evolution include logarithmic and convergent behavior, while changes in population size due to positive selection for fecundity become less pronounced. This two-phase evolutionary trajectory does not seem to be strain-dependent, and varies from strain to strain only in terms of its absolute timing and magnitude.

Both of these phenomena (dual effects of positive selection and two-phased evolutionary trajectory) can be explained in terms of fitness landscape dynamics. An empirical fitness landscape, or genotype-fitness map [13], provides us with a surface to characterize both fitness gains (hill-climbing) and losses (valley descent). In terms of characterizing trends in population size amongst our genealogies, populations representing those strains that demonstrate a clear logarithmic increase in fecundity over evolutionary time have climbed towards a fitness peak. By contrast, populations representing strains that demonstrate fluctuation in fecundity over evolutionary time are traveling the neutral parts of the fitness landscape and even exploring local fitness minima. For some genetic backgrounds, selecting for fecundity is not particularly advantageous from both a developmental and reproductive standpoint. This is consistent with Stoltzfus [14], who argues that an evolving population evolves towards a fitness peak via mutational diversity rather than the optimization of traits.

The question remains as to what could be driving these processes at the genetic level. Recall that we observe generalized patterns that do not directly corresponding to mutations on specific genes or gene families. This suggests the products of selection are due to antagonistic pleiotropy [15] and other complex interactions [16]. Even though we are studying hermaphroditic genetic mutant lines subject to extensive backcrossing, we are still generating thousands of random mutations per strain. While these mutations may be neutral and occur at very low frequencies, but they may also contribute to individual variation and greater evolvability. For a better understanding of this, we need to link systematic gains and losses in fecundity to their potential life-history mechanisms. One the one hand, some mutant genotypes recapitulate the fitness maximization patterns of the wildtype, albeit at a lower order of magnitude. In the case of our AMPK mutants, this effect is due to the characterized function of these mutants, which is not a developmental defect *per se*, but rather a physiological defect which may impair their reproductive capacity. On the other hand, the *lag* and *daf-c* mutants exhibit specific defects in developmental processes. While the function of these mutants are not directly tied to changes in the heterochronic timing [17, 18] of reproduction, they might nevertheless act against fitness gains that would otherwise result from positive selection. This latter point is consistent with Lang and Desai [19] who argue that experimental evolution tends to produce fitness increases via epistasis and parallel pathways associated with diverse sets of mutations.

## ACKNOWLEDGEMENTS

We would like to thank Dr. Nathan Schroeder, Rebecca Androwski, Kristen Flatt for their advice on *C. elegans* classical genetics techniques and worm culture. I would also like to thank viewers of poster 766C at the 20^th^ International *C. elegans* Meeting for their comments. A portion of the work presented in this paper was supported by the National Institute of General Medical Sciences of the National Institutes of Health (Award Number IR01GM111566-01). Thanks also go to the experimental evolution community, which also served as inspiration for this study. Some strains were provided by the CGC, which is funded by NIH Office of Research Infrastructure Programs (P40 OD010440).

## METHODS

All data (genealogies, control conditions) and selected measures (RCO) can be found on Figshare at doi:10.6084/m9.figshare.2087719.

### Organisms

**Strains:** Seven (7) strains were acquired from either the Caenorhabditis Genomics Center (CGC, http://cbs.umn.edu/cgc/) or Nathan Schroeder’s Laboratory at University of Illinois Urbana-Champaign. These genotypes were: qIs56[Plag-2::GFP]; daf-7(e1372), qIs56[Plag-2::GFP]; daf-9(rh50), N2, myIs14[Pklp-6::GFP]; daf-7(e1372); daf-16(mu86), qIs56[Plag-2::GFP], lag-2(q420), and daf-16(mu86). Further information regarding the [Plag-2::GFP] and [Pklp-6::GFP] constructs can be found in [20].

**Genetic Crosses:** Seven (7) additional crosses were constructed to gain additional data about the contributions of the AMPK and daf-7 mutant genotypes to evolvability and developmental plasticity. These genotypes were: qIs56[Plag-2::GFP]; daf-7(e1372); daf-16(m27), qIs56[Plag-2::GFP]; aak-1(tm1944); daf-7(e1372), qIs56[Plag-2::GFP]; aak-2(ok524); daf-7(e1372), qIs56[Plag-2::GFP]; aak-1(tm1944); aak-2(ok524); daf-7(e1372), qIs56[Plag-2::GFP]; aak-1(tm1944), qIs56[Plag-2:: GFP]; aak-2(ok524), qIs56[Plag-2::GFP]; aak-1(tm1944); aak-2(ok524). More details on the design can be found on Figshare at doi: 10.6084/m9.figshare.2436289.v1

### Measures

**Reproductive Carry-Over.** Reproductive Carry-Over (RCO) is a discrete, subtractive first-order time derivative of the population size. RCO can be calculated recursively on an averaged time-series, or on individual genealogies. RCO is formally stated as

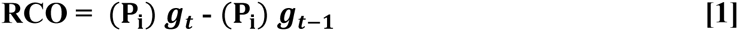

where *P*_i_, are populations derived from the same founder worm, *gt* is the current generation in the genealogy, and *gt-1* is the previous generation in the genealogy. This results in 19 timepoints for an evolutionary trajectory of 20 generations.

**Exponential Smoothing.** Exponential smoothing is used to analyze the population size time-series by removing the noise of demographic fluctuation and setting the initial condition of the smoothed Generation 1 to a value of 1. The exponential smoothing kernel is applied recursively and defined as

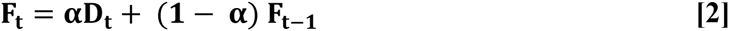

where *F*_*t*_ is the current point in the smoothed evolutionary trajectory, *D*_*t*_ is the corresponding timepoint in the unsmoothed time-series (population size), *F*_*t*-1_ is the previous point in the smoothed evolutionary trajectory, and *α* is the smoothing constant.

**Normalized Population Size.** Normalized Population Size is the measured population size for a single genealogy or population average in a single generation subtracted from a control population size for a given genetic strain derived using the hanging drop method. Normalized population size is calculated for every generation in the evolutionary trajectory against a constant value for the control population.

### Statistics and Analyses

**Graphs.** All graphs and data presentation were made in Excel, MATLAB, and R. All statistical analyses were run in MATLAB and R.

### Techniques

**Determining Population Size.** For every generation, five replicates were created. After a 60mm diameter plate filled with NGM agar is seeded with 100uL of OP50 media and allowed to dry, a single L4 stage worm is placed on each of the five plates. Each plate is allowed to grow for 3d at optimal temperatures for the given strain. At the end of the 3d period, plates are examined for offspring and counted. Anywhere from 1-3 plates then are selected to represent the subsequent generation. Five new plates are seeded from the selected populations with L4 stage representatives in proportion to their population size (e.g. larger populations get more representatives in the next generation).

**Selection criterion.** As discussed in the population size section, all populations were selected for fecundity at 3d after being derived from L4 stage individuals. In cases where no offspring were produced, the window for selection was increased to 6d based on observations of a delayed life-history. Populations were selected on the basis of fecundity and their population size relative to the other replicates populations in that observation. Therefore, the selection criterion was not absolute, but was consistently applied across all strains.

**Hanging Drop method.** The hanging drop method was proposed by [21] as a means of assessing the fecundity of a single worm over a discrete period of time isolated from maternal effects and with minimal counting error. The hanging drop is conducted by harvesting a series of worms at the L4 stage and plating them one worm per plate. Every 24h, each plate is checked for offspring. The number of offspring are counted, and the parent is transferred to a new plate. This was done over the course of 3d for each evolved strain to serve as a control. The population count for each strain was derived by summing all three daily measurement per replicate and then averaging across the replicates.

**Genotyping for Mutant Construction.** Genotyping for the aak-1(tm1944) and aak-2(ok524) mutants were done using the following primers: tm1944; internal (TCACACGTCTCTTCCGTGTT),left flanking (TCGCGTCCAGAAGAAGATTT), right flanking (TCCCTTTCTTCGCTCACTTT). ok524; internal (CAAAGTCCGCAATCTTCACA), left flanking (TCATCCGCCTCTACCAAGTC), right flanking (TCAAATCCCATTTCGCTTTC). Sequences for primer design retrieved from Genbank http://www.ncbi.nlm.nih.gov/. Primer design conducted using Primer3 (http://biotools.umassmed.edu/primer3/primer3web_input.htm). Using Blastn, primers were evaluated (e.g. E-value) for similarity with bacterial sequences and other Nematode sequences.

